# Bayesian Hierarchical Hypothesis Testing in Large-Scale Genome-Wide Association Analysis

**DOI:** 10.1101/2024.02.26.582204

**Authors:** Anirban Samaddar, Tapabrata Maiti, Gustavo de los Campos

## Abstract

Variable selection and large-scale hypothesis testing are techniques commonly used to analyze high-dimensional genomic data. Despite recent advances in theory and methodology, variable selection and inference with highly collinear features remain challenging. For instance, collinearity poses a great challenge in Genome-Wide Association Studies (GWAS) involving millions of variants, many of which may be in high linkage disequilibrium. In such settings, collinearity can significantly reduce the power of variable selection methods to identify individual variants associated with an outcome. To address such challenges, we developed a Bayesian Hierarchical Hypothesis Testing (BHHT)–a novel multi-resolution testing procedure that offers high power with adequate error control and fine-mapping resolution. We demonstrate through simulations that the proposed methodology has a power-FDR performance that is competitive with (and in many scenarios better than) state-of-the-art methods. Finally, we demonstrate the feasibility of using the proposed methodology with big data to map risk variants for serum urate using data (n∼300,000) on phenotype and ultra-high-dimensional genotypes (∼15 million SNPs) from the UK-Biobank. Our results show that the proposed methodology leads to many more discoveries than those obtained using traditional feature-centered inference procedures. The article is accompanied by open-source software that implements the methods described in this study using algorithms that scale to biobank-size ultra-high-dimensional data.

---

Many modern statistical learning problems require detecting from a large number of features a small fraction of them that are jointly associated with an outcome. In recent years, many variable selection methods have been proposed (for reviews see e.g., Fan and Lv (2010), O’Hara and Sillanpää (2009), Lu and Lou (2022)). One can also pose a variable selection task as a multiple hypothesis testing problem. This approach uses decision-theory to optimally balance power and type-I errors.

Despite the important advancements in theory and methods for high-dimensional regression, variable selection and inference in the presence of highly collinear features remain challenging Efron (2010). The problem of selecting a subset of predictors among a large set of correlated features is ubiquitous in Genome-Wide Association (GWA) studies where the objective is to map regions of the genome (either individual variants or chromosome segments) associated with a phenotype. In the last decade, several public (e.g., UK-Biobank, Million Veteran Program, TOPMed, All of Us) and private (e.g., 23andMe®) initiatives have generated unprecedentedly large biomedical datasets comprising genotypes linked to extensive phenotype and disease information. The advent of big data brings unprecedented opportunities to advance genetic research and predict complex traits (de los Campos *et al*. (2018)). However, together with larger sample sizes, these modern datasets come with a remarkable increase in marker density, with potentially millions of single nucleotide polymorphisms (SNPs) distributed throughout the genome. With such a high SNP density, many SNPs can be highly correlated due to linkage disequilibrium (LD).

Bayesian variable selection methods (BVS) (George and McCulloch (1997), Ishwaran and Rao (2005)) can be used to identify risk variants in GWA studies. Following the seminal contribution of Meuwissen *et al*. (2001), these methods have been widely used for the prediction of complex traits in plant and animal breeding (e.g., Meuwissen *et al*. (2001), de los Campos *et al*. (2009), Habier *et al*. (2011)) and also in human genetics (e.g., de los Campos *et al*. (2010), Makowsky *et al*. (2011), Guan and Stephens (2011), Kim *et al*. (2017), Lello *et al*. (2018), Lee *et al*. (2018), Wang *et al*. (2020)).

Bayesian methods offer adequate quantification of uncertainty in variable selection problems. This feature (which is essential for any inferential task) can have unwanted consequences when the goal is to select individual variants associated with a phenotype. For example, when multiple variants are all in LD with one or more causal variants, the posterior probability of non-null effect (aka posterior inclusion probabilities or PIPs) of individual SNPs may not achieve a high value for any individual locus because of the uncertainty about individual SNP effects introduced by collinearity.

Therefore, variant-centered BVS may not identify important risk loci when those loci are located in regions of high LD. This phenomenon has been reported and studied before (Ghosh and Ghattas (2015)) and the general recommendation is to avoid using marginal posterior probabilities and focus instead on credible set inferences, i.e., identifying sets of predictors that are jointly associated with an outcome with a high level of credibility. The joint posterior distribution of BVS contains all the information needed to identify such sets. However, we lack methods to identify credible sets from posterior samples efficiently.

We fill this gap by developing a methodology for multi-resolution Bayesian hypothesis testing that can lead to powerful inferences, with adequate error control, and fine-mapping resolution. Our approach integrates ideas first proposed for frequentist hierarchical hypothesis testing (Yekutieli (2008),Meuwissen *et al*. (2001), Barber and Ramdas (2016), Renaux *et al*. (2020), Sesia *et al*. (2020)) into a Bayesian framework that can offer powerful inferences with adequate error control.

Our study makes the following contributions: (i) We develop an algorithm to identify credible sets from posterior samples. (ii) Using simulations, we show that the proposed algorithm can achieve high power with low error rate and fine mapping resolution, and perform similarly (and in some scenarios better) than stat-of-the-art credible set inference methods. Further, (iii) We demonstrate the feasibility of using the proposed methods with biobank-size data and ultra-high-density SNP genotypes. Finally, (iv) We provide open-source software that implements the proposed algorithms.

## Materials and Methods

Consider a multiple regression model of the form,

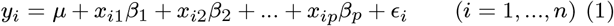

where *y*_*i*_ is a phenotype of interest, *x*_*i*1_, …, *x*_*ip*_, are predictors (e.g., SNPs or features from other omics), *β*_1_, …, *β*_*p*_ are individual feature effects, and *ϵ*_*i*_ is an error term for the i-th individual in the sample of size *n*.

Bayesian Variable Selection models often use IID priors from the Spike-and-Slab family (George and McCulloch (1993), Ishwaran and Rao (2005)), which include a point of mass at zero (i.e., no-effect, *β*_*j*_ = 0) and a slab (non-zero effect) of the form

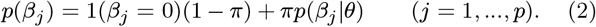

Above, *π* is the prior probability of a non-zero effect, *p*(*β*_*j*_|*θ*) is a density function (the ‘slab’) describing the distribution of non-null effects (e.g., a Gaussian prior), and *{*1(*β*_*j*_ = 0) = 1 *if β*_*j*_ = 0; 0 *otherwise}* is an indicator function.

The samples from the posterior distribution of BVS models can be used to estimate the (marginal) posterior probability of non-zero effects for each of the predictors:

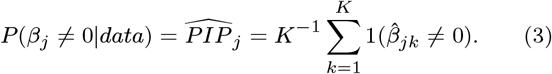

where *k* = 1, …, *K* is an index for posterior samples 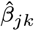 for *j* = 1, …, *p*. These posterior probabilities can be used to identify predictors that are confidently associated with an outcome, e.g., those with 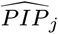 *≥* 0.9 corresponding to a local-False Discovery Rate (LFDR) *≤* 0.1. However, as noted earlier, marginal inferences can be under-powered in the presence of collinearity.

### Credible Set Inference

A level-*τ* (*τ ∈*[0, 1]) credible set is a set of predictors whose joint probability of association is greater that *τ* (Wang *et al*. (2020)). The joint probability of association of a set is the probability that at least one of the features in the set is associated with the outcome. For a set of predictors Ω, the set-PIP (denoted by *PIP*_Ω_) can be defined as follows.

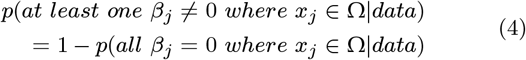

These set-PIPs can be estimated using samples from the posterior distribution by simply counting the proportion of posterior samples that contain at least one of the effects in the set different than zero, for a set Ω = *{j, j*^*′*^*}*:

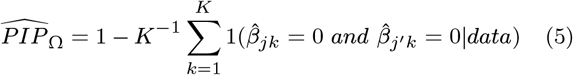

Next, we introduce a Bayesian Hierarchical Hypothesis Testing (BHHT) algorithm to identify *α*-level credible sets.

### Bayesian Hierarchical Hypothesis Testing

(BHHT) tests a sequence of nested hypotheses associated with hierarchical clusters of the inputs (*x*_*i*1_, …, *x*_*ip*_). The procedure is schematically described in Fig. 1 which depicts an example involving four features clustered according to a binary tree. Testing starts at the top node; every time a hypothesis is rejected, the two nested hypotheses involving the effects under the left and right branches are tested. In Fig. 1, nodes with red (white) squares denote null hypotheses that were (were not) rejected. The path containing rejected nulls is marked in red. In the example in Fig. 1, testing involves four steps:

1. *H*_(0(1))_ : *β*_1_ = *β*_2_ = *β*_3_ = *β*_4_ = 0 Vs *H*_(*a*(1))_ : at least one coefficient different than 0. Decision: reject *H*_(0(1))_.
2. *H*_(0(2*L*))_ : *β*_1_ = 0 Vs *H*_(*a*(2*L*))_ : *β*_1_ ≠ 0, and *H*_(0(2*R*))_ : *β*_2_ = *β*_3_ = *β*_4_ = 0 Vs *H*_(*a*(2*R*))_ : at least one of the three coefficients ≠ 0. Decisions: Do not reject *H*_(0(2*L*))_ and reject *H*_(0(2*R*))_.
3. *H*_(0(3*L*))_ : *β*_2_ = 0 Vs *H*_(*a*(3*L*))_ : *β*_2_ ≠ 0, and *H*_(0(3*R*))_ : *β*_3_ = *β*_4_ = 0 Vs *H*_(*a*(3*R*))_: either *β*_3_ or *β*_4_ or both coefficients ≠ 0. Decisions: reject both *H*_(0(3*L*))_ and *H*_(0(3*R*))_.
4. *H*_(0(4*L*))_ : *β*_3_ = 0 Vs *H*_(*a*(4*L*))_ : *β*_3_ ≠ 0 and *H*_(0(4*R*))_ : *β*_4_ = 0 Vs *H*_(*a*(4*R*))_ : *β*_4_*≠*0. Decision: no null rejected.

**Figure 1.**
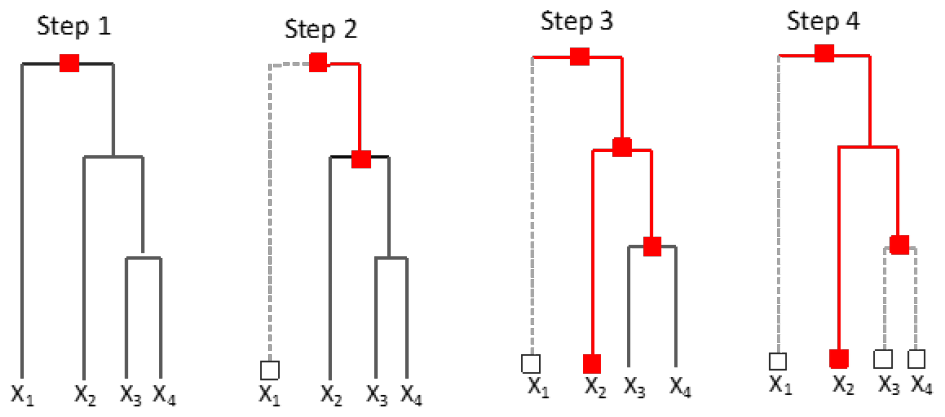
Schematic representation of Hierarchical Hypothesis Testing. The example involves 4 features (*X*_1_−*X*_4_). Testing starts at the top node and proceeds to lower levels of the hierarchy, whenever a null hypothesis at the parent node is rejected (red squares). If a null hypothesis is not rejected (white squares) no further testing is performed within the branch.

The final discovery set includes two elements *X*_2_ (a singleton) and a duplet *{X*_3_, *X*_4_*}* .

### Error control

We focus on controlling the Bayesian FDR of the discovery set (BFDR, controlling BFDR at an *α*-level also provides FDR control at the same level, see Lemma S1 in the Appendix).

We propose an algorithm for discovery set finding which control the DS-BFDR in BHHT. The algorithm conducts a sequence of tests over a grid of values of thresholds for the LFDR’s 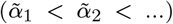. For each value in the grid, we determine the 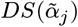 and estimate the DS-BFDR using (5). If the estimated DS-BFDR is below the target threshold (e.g., *α <* 0.05) we move to the next *α* in the grid, otherwise, if DS-BFDR *≥α*, we use 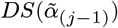 as the final DS. For the grid of threshold values 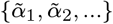, we use the ordered LFDRs from the tree since the DS-BFDR as a function of the thresholds changes only at these values. Algorithm S2 of the Appendix provides further details about the proposed methodology.

The function `Bayes_HHT` included in the GitHub repository associated with this manuscript (see Data availability section) takes as inputs posterior samples, a hierarchical clustering of predictors (i.e., a ‘merge’ object) and a BFDR *α*-value and returns credible sets obtained through BHHT. By default, this function performs BHHT controlling the BFDR; however, we also offer the option to control the node FDR as well as the Sub-family FDR Yekutieli (2008) (see appendix for further details).

### Demonstration with a simplified example

To demonstrate the application of BHHT we present a simplified example involving mice genomic data from Wellcome Trust Valdar *et al*. (2006). The data used in the example consists of genotypes of 1814 mice. We used the first *p* = 300 SNPs from chromosome 1 to simulate a phenotype (*y*) under the linear regression model (Eq. 1) with Gaussian iid error terms. Only four of the 300 SNPs had non-zero effects. We fixed the error variance to ensure that the four causal variants explained 10% variance of the phenotype (*y*). Subsequently, we used the BGLR R-package Pérez and de los Campos (2014) to generate samples from the posterior distribution of a Bayesian model with a spike-slab prior (2). The code used to perform this simulation is available in a GitHub repository linked to the paper (see Data Availability section).

Fig. 2 shows the SNP-specific posterior probability of non-zero effects (PIPs). No individual SNP had a PIP>0.95 (dashed horizontal line, corresponding to a local BFDR of 0.05). However, a BHHT algorithm (controlling the DS-BFDR at *α* = 0.05) successfully identified four sets (Table 1) with high set-PIP each containing a causal variant (see vertical shaded area in Fig. 2 and Table 1). Note that, the second credible set in Table 1 has local-FDR (1-Set-PIP=0.09) greater than 0.05 and would have been excluded from the discovery set by a local (node or Sub-family FDR) FDR controlling method. However, this set is included as the BFDR of the discovery set (DS-BFDR = 1 *−*0.955 = 0.045) is below 0.05. This demonstrates the gain in power by controlling the discovery set BFDR compared to the local-FDR controlling strategies.

**Table 1:** Four credible sets discovered through Bayesian Hierarchical Hypothesis Testing using the node BFDR criteria (variants with non–zero effect in the simulation are highlighted in bold).

| Features included | Set size | Set-PIP | Local BFDR |
| --- | --- | --- | --- |
| 63, 64, 65, **66**, 68 | 5 | 0.95 | 0.05 |
| **125**, 126, 129 | 3 | 0.91 | 0.09 |
| 221, 223, 218, **219**, 220, 222, 216, 217, 224, 225, 226, 228, 227 | 13 | 0.96 | 0.04 |
| 270, 272, 271, 266, **267** | 5 | 1.00 | 0.00 |
| **Entire Discovery Set** | **26** | **0.955** | **0.045** |

**Figure 2.**
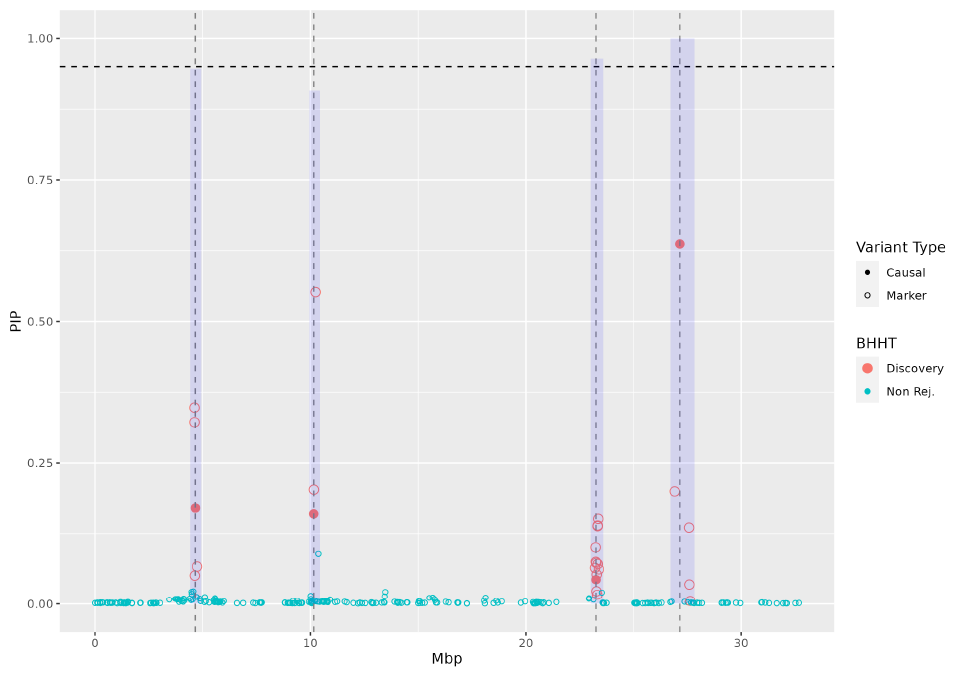
Posterior Probability of non-zero effect by SNP and Credible Set (CS) identification using Bayesian Hierarchical Hypothesis Testing (BHHT). Each point represents a SNP, the vertical axis gives the posterior probability of non-zero effect (PIP) and the horizontal axis is the map position in mega-base-pairs (Mbp). The purple vertical bars give the joint posterior probability of association of each of the CS identified through BHHT (see Table 1 below).

### Simulation studies

We used simulations to benchmark BHHT against inferences based on marginal PIP and against SuSiE (Wang *et al*. (2020)) – a state-of-the-art procedure for CS inference proposed for GWA studies. We perform two simulation studies, one where we simulated predictors and the outcome (this setting allowed us to control the level of collinearity) and one using real human genotypes with biobank-like sample size and ultra-high-density genotypes.

#### Simulation 1

For each of the 1000 Monte Carlo replicates we simulated 525 SNPs using a Markov process such that the correlation between predictors was 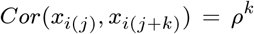. Here, *i* is an index for the subject, *j* is an index for the feature in the sequence, and *k* is the lag-between predictors. We considered two scenarios for collinearity: *ρ* = 0.45 (moderate) and *ρ* = 0.90 (high). Then, we simulated the response (*y*_*i*_) according to the linear model (Eq. 1) with either 5 or 20 (*S* = 5 or 20) of the 525 predictors having non-zero effects. The errors were *iid* Gaussian with zero mean and a variance tuned such that the proportion of variance of the outcome explained by the signal (aka heritability, *h*^2^) was 0.0125. Further details about the simulation can be found in the Appendix.

#### Simulation 2

For this simulation we used real genotypes from the UK Biobank and and a phenotype simulated from those genotypes. In this case, to vary the level of collinearity among predictors we consider two scenarios: (2.1) SNPs from the arrays used in the UK Biobank (aka calls, *∼*785,000 SNPs) and (2.2) SNPs from imputed genotypes provided by the UK-Biobank (the imputed SNPs were filtered to satisfy minor-allele-frequency *≥*0.1 % and missing call rate *≤*5 %, 15 millon SNPs). The calls dataset contains SNPs that are physically distant representing scenarios with low correlation whereas the imputed dataset contains highly dense SNPs exhibiting LD blocks with near-perfect collinearity.

For each of the 500 Monte Carlo replicates, we randomly sample a chromosome and, within it, a 100 kilobase pairs block of SNPs. Within each block, we sampled five loci and used them as the causal variants to generate a phenotype using a linear model (Eq. 1) with Gaussian errors and a segment-heritability (*h*^2^) of 0.0125. When fitting models, we used not only all the SNPs in the 100 kbp holding the causal loci but also those in the 500 kbp flanking regions.

## Statistical Analysis

We analyzed the simulated data using a Bayesian linear model with IID spike-slab priors (described in Eq. 2) for the regression coefficients. Models were fitted using the BGLR R-package Pérez and de los Campos (2014). Then, we performed hierarchical clustering (Hartigan (1975) Ward Jr. (1963)) of the SNPs using the R function hclust and applied BHHT. We considered two benchmarks for BHHT:

i. **Marginal hypothesis testing:** We used marginal SNP-PIPs to produce discovery sets (see Algorithm S3 of the Appendix for further details), and
ii. **SuSiE:** A credible-set variable selection procedure proposed in Wang *et al*. (2020). This method uses a prior distribution that leads to the formation of credible sets. We fitted SuSiE and obtained the credible sets using the R-package susieR Wang *et al*. (2020).

For all methods, we vary the BFDR threshold *α* in the set *{*0, 0.02, 0.05, 0.1*}*. For the detailed implementation please refer to the code repository in the Data Availability section.

### Real data analysis

To demonstrate the feasibility of using BHHT with large-scale GWAS data, we applied BHHT to data from the UK Biobank dataset (*n ∼*300, 000 unrelated individuals of European ancestry), using serum urate as the phenotype and SNP genotypes from either (1) the SNP array or (2) from the imputed genotypes. We first log-transform the serum urate levels to make it symmetrically distributed and then adjust it with respect to age, sex, center, and top 10 SNP-derived principal components.

Due to the high SNP-density, the imputed genotypes have LD blocks with near-perfect collinearity; therefore the analyses involving the ultra-high-density SNPs serve as an example of how the proposed methods can work under extreme collinearity.

It is computationally infeasible to fit a multiple regression model jointly with 15 million SNPs. However, in the white European UK-Biobank cohort LD decays rather quickly with physical distance getting to values very close to zero at 500 kilobase pairs to 1 megabase pairs. Therefore, following Funkhouser *et al*. (2020), we conducted the association study by fitting local Bayesian regression models to overlapping windows containing 9000 SNPs which were displaced by 1000 SNPs; thus, the segments analyzed had a core of 7000 SNPs and 1000 SNPs in each of the flanking regions. For each window, we fitted models to the 9000 SNPs but retrieved the posterior samples of the SNPs in the core (7000 SNPs) only. We first selected the set of individual SNPs with BFDR less than 0.05 (Algorithm S3) and group the selected SNPs into segments consisting of all SNPs with PIP>0.05 that were at a distance smaller than 100 Kbp. Then, we applied BHHT within each of these segments. Additional details of the analysis strategy are discussed in the Appendix.

## Results

### Simulation 1

#### Power-FDR performance

Fig. 3 shows the power-FDR performances of SNP-PIP, BHHT, and SuSiE when the discovery sets were restricted to have at most 5 predictors. In this setting, when the number of causal effects was equal to 5 and collinearity was moderate (top-left panel in Fig. 3) we observed that all methods performed similarly, with the method based on marginal probabilities (SNP-PIP) performing only slightly worse in the scenario with *n*=10K. However, for the same trait architecture, when collinearity was high (*ρ* = 0.9 the bottom row of the left panel) the two credible set methods (DS-BFDR and SuSiE) outperformed marginal hypothesis testing (SNP-PIP) highlighting the power of credible set inference. In the scenario with only 5 causal variants and high collinearity, SuSiE performed slightly better than BHHT. However, when we considered a model with more causal variants (20 predictors with non-zero effect, panel B in Fig. 3) BHHT outperformed SuSiE. This may reflect a limitation of the variational algorithm in SuSiE which in multi-modal problems can get stuck at local optima (see the Discussion of Wang *et al*. (2020)). Regardless of the scenario and method, as one would expect, for any FDR level, power was higher with the largest sample size.

**Figure 3.**
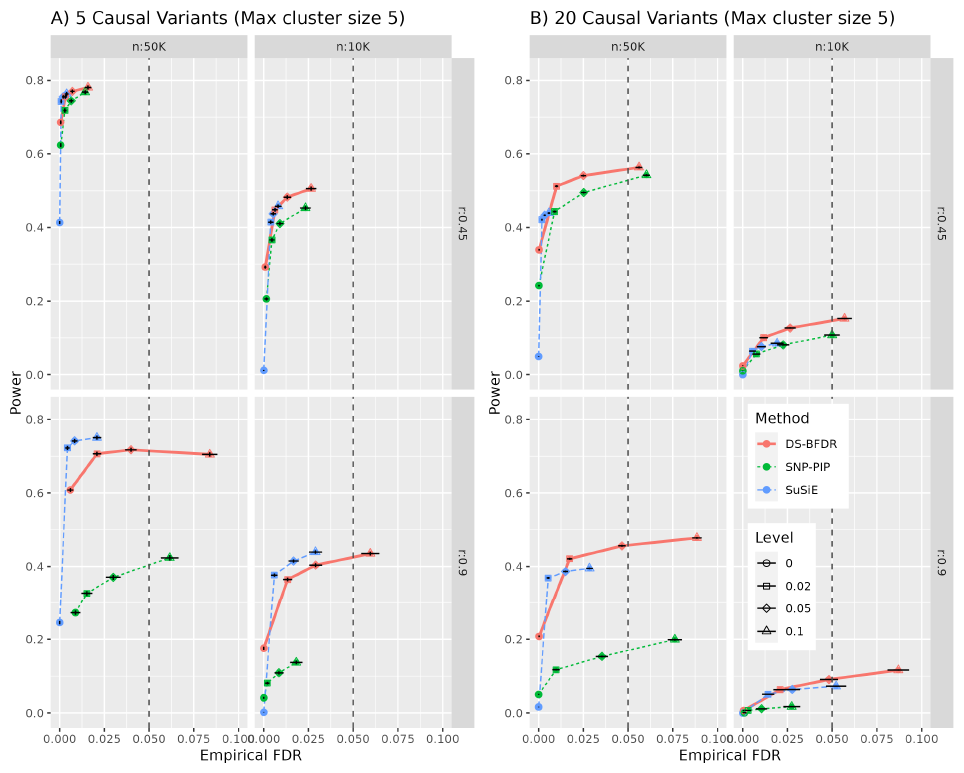
Power vs FDR curves for <u>simulation one</u>, by simulation setting. Here, *α, n*, and *r* indicate the BFDR threshold used, sample size, and the lag-1 correlation of the simulated predictors, respectively. The Left and right panels represent scenarios where there are 5 and 20 causal variants respectively. In all scenarios, predictors explained 1.25% of the variance of the outcome. As collinearity increases the credible set inference gains significant power over individual SNP-level inference. Furthermore, the proposed method achieves higher power than SuSiE when there exists a greater number of causal variants in the true model.

We also performed the same power-FDR evaluation restricting the maximum credible set size to 3 or 10 variants. The results from these analyses are presented in Fig. S2 and Fig. S3, respectively. The overall results were conceptually similar to the ones presented in Fig. 3 with a few differences: (i) Imposing a more stringent restriction on the maximum cluster size reduced the power advantage of the CS inference methods relative to marginal SNP-PIP inferences (ii) When the maximum CS size was restricted to be only 3 SNPs, there were small differences between SuSiE and BHHT, and, finally, (iii) when the maximum CS size was allowed to be 10 SNPs, BHHT outperformed SuSiE by a sizable margin in the scenario with high LD and 20 causal variants.

### FDR control

Previously, we observed that DS-BFDR can achieve high power at low FDR even in the scenario of high collinearity. However, for a decision rule to be useful it must provide adequate error control. In each panel of Fig. 4, we show the empirical FDR vs the BFDR threshold *α* for the four simulation settings. For all settings, we observed that all methods accurately controlled the FDR at the desired level.

**Figure 4.**
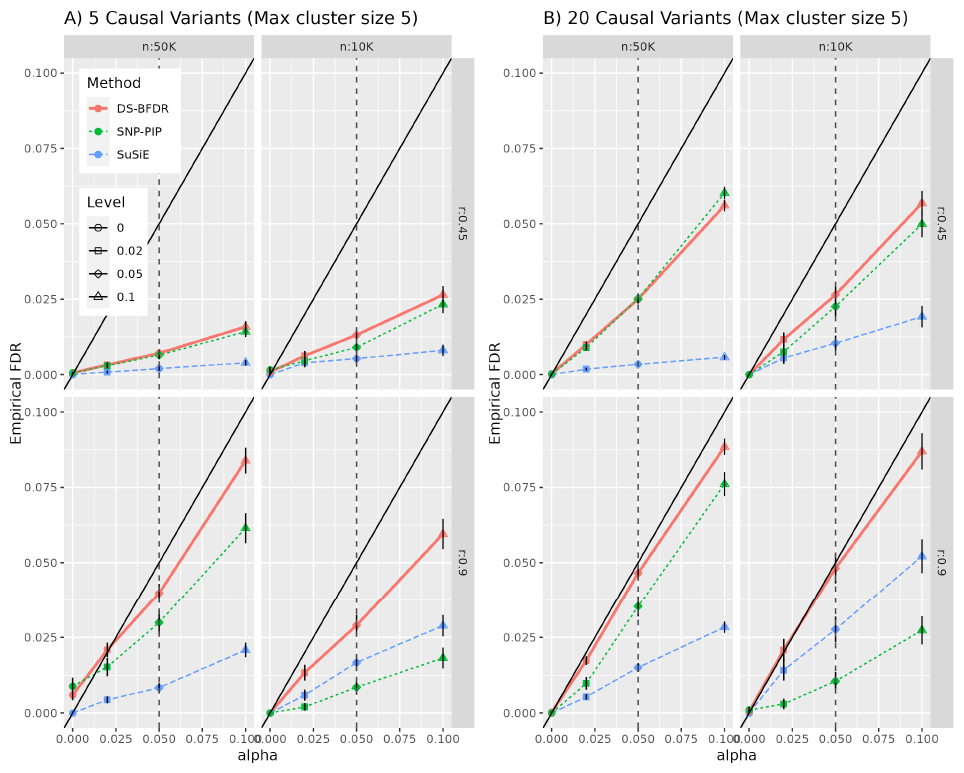
Empirical False Discovery Rate (FDR) vs level *α* plot for <u>simulation one</u>, by simulation setting. Here, *α, n*, and *r* indicate the BFDR threshold used, sample size, and the lag-1 correlation of the simulated predictors, respectively. The Left and right panels represent scenarios where there are 5 and 20 causal variants respectively. In all scenarios, predictors explained 1.25% of the variance of the outcome. All the methods accurately control the FDR.

### Simulation 2 (real genotypes)

Building on the simulation results of the previous section, we evaluated the BHHT (and the benchmark methods) using phenotypes simulated from real genotypes from UK-Biobank. To evaluate the power-FDR performances of the methods we construct the simulation scenarios by varying: (i) **degrees of collinearity** (calls or imputed data), (ii) **sample sizes** (n=10K,100K, and 300K, K=1,000), and (iv) **mapping resolution**. For the latest, following de Los Campos *et al*. (2023), we defined a SNP in a credible set as a true discovery if the the mapping distance from the SNP to the closest causal variant was smaller than *x* kilobase pair (kbp). To compare methods at different mapping resolutions, we varied *x* from 0kbp (i.e., perfect mapping resolution) to 10kbp.

### Power-FDR performance

Fig. 5 shows the power-FDR curves for SuSiE, SNP-PIP, and BHHT by SNP set (calls and imputed, in columns) and sample size (in rows). The results in Fig. 5 are based on a perfect mapping resolution (*x* = 0kbp). When the correlation among the SNPs was moderately low (calls dataset in the left column) all the methods had similar power with very small FDR (especially for large sample sizes). However, with the imputed dataset (which has highly correlated SNPs, right column in Fig. 5), the two credible set methods (DS-BFDR, and SuSiE) have significantly higher power than the marginal hypothesis testing (SNP-PIP). In addition, we observed that the DS-BFDR attains similar levels of power as SuSiE with significantly lower FDR levels. This might be because SuSiE is prone to discover credible sets with high FDR (see Fig. 3 in Wang *et al*. (2020)). Across all methods on the two datasets, we observe that the maximum power has been attained by the largest sample size scenario.

**Figure 5.**
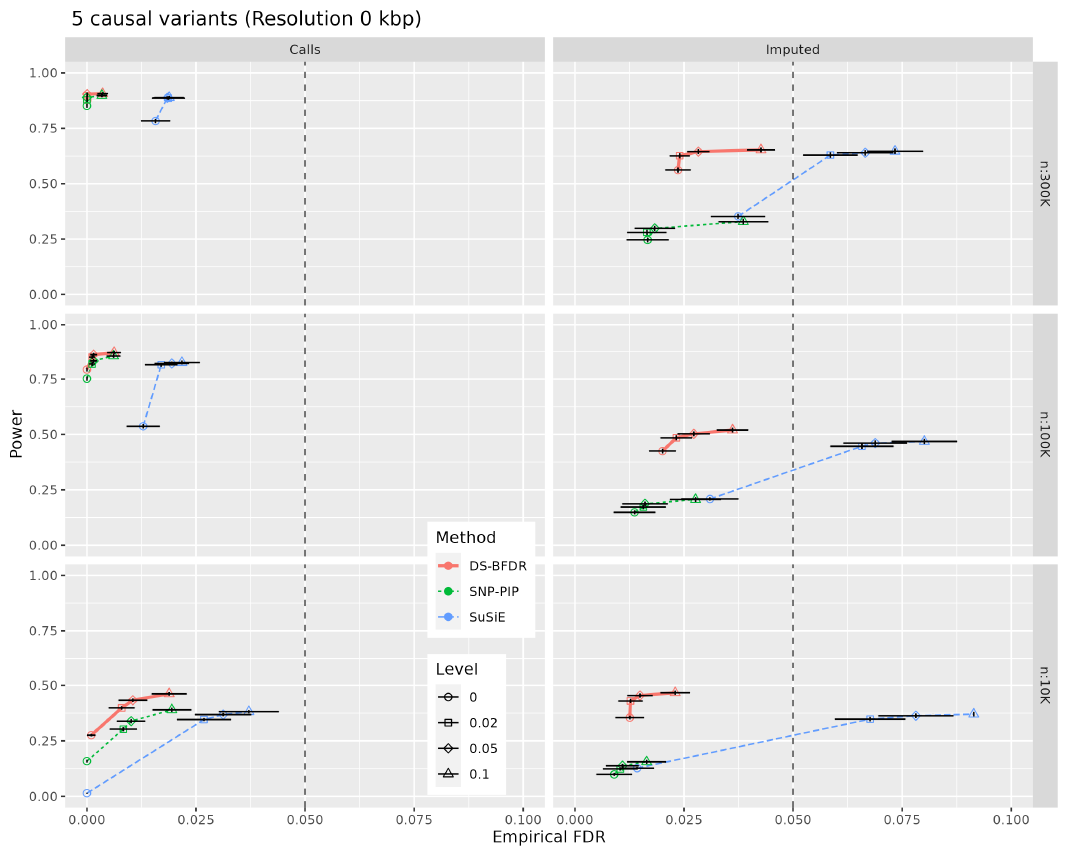
Power vs FDR curves for simulation two by sample size (rows) and the SNP set used (columns). Here, *α* and *n* indicate the BFDR threshold used and sample size respectively. In all scenarios, SNPs explained 1.25% of the variance of the outcome. Credible set inference gains significant power over individual SNP-level inference in the presence of strong collinearity (imputed SNPs). With the imputed dataset, Bayesian Hierarchical Hypothesis Testing (BHTT) achieved similar power as SuSiE with a significantly lower FDR.

For a 10 kbp mapping resolution, the power-FDR performances of the methods (shown in Fig. S5) were similar to those in Fig. 5 except that we observed a significant decrease in FDR for all methods which happens because some discoveries that are deemed false for a perfect mapping resolution (*x* = 0kbp) are considered true with a 10kbp tolerance.

### Large-Scale GWAS Application with Real Phenotype

In the real data analyses, we found 123 SNPs with significant individual SNP-PIP (i.e., BFDR of the set *≤*0.05) in the analysis using the calls SNPs and only 98 SNPs satisfying this criterion in the analysis using imputed SNPs (Table 2). The fact that fewer findings were obtained with the imputed SNPs (which include the genotyped SNPs as a subset) may seem counter-intuitive. However, this happened because LD is much stronger in the imputed SNPs than in the calls. Under these conditions, due to high LD, many SNP that are in regions harboring causal variants do not reach high SNP-specific PIP.

**Table 2.**
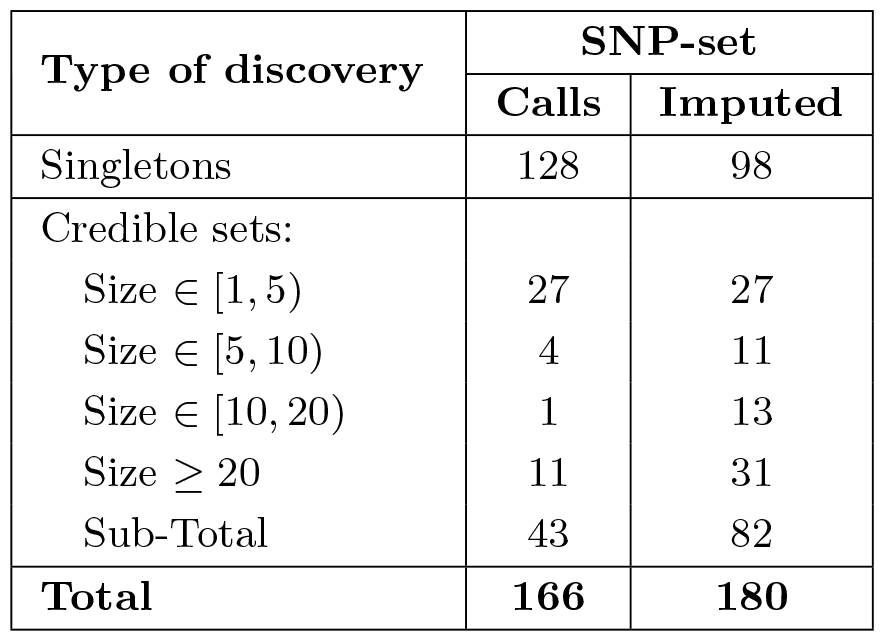
Number of discoveries for serum urate by SNP-set used and type. The singletons and credible sets are from applying the marginal testing and BHHT algorithm respectively with *α* = 0.05.

| Type of discovery | SNP-set |  |
| --- | --- | --- |
|  | Calls | Imputed |
| Singletons | 128 | 98 |
| Credible sets: |  |  |
| Size $\in [1, 5)$ | 27 | 27 |
| Size $\in [5, 10)$ | 4 | 11 |
| Size $\in [10, 20)$ | 1 | 13 |
| Size $\geq 20$ | 11 | 31 |
| Sub-Total | 43 | 82 |
| **Total** | **166** | **180** |

However, using BHHT applied to segments harboring SNPs with elevated PIP yielded 43 additional discoveries (with DS BFDR *≤*0.05) for the calls and 82 for the imputed genotypes; thus making the total number of credible sets discovered to be 166 with the calls and 180 when using the imputed genotypes. Thus, the use of CS inference increased the number of discoveries by 35% and by 83% for the calls and imputed genotypes, respectively (Table 2). These results highlight the importance of using credible set inferences when using ultra-high-density genotypes. Most of the credible sets identified through BHHT were small (between 1 to 4 SNPs); however, some credible sets involved more than 20 SNPs (Table 2). The physical positions of the credible sets (CS) identified (using the imputed genotypes) through BHHT are displayed in an ideogram in Fig. S7.

To provide further insight on how BHHT works, we showcase in Fig. 6 a plot of the SNP-PIPs of a segment in chromosome 6 which contains a credible set (found through BHHT) formed by four SNPs (denoted with solid red points in the top PIP plot in panel a) which, individually, had low PIP (*∼*0.3). These SNPs won’t be identified as significantly associated with serum urate when inferences are based on individual SNP PIPs. However, jointly, the four SNPs achieve a high set-PIP. The LD-heatmap displayed in panel (b) of Fig. 6 shows that the five SNPs in the credible set are in high mutual LD.

**Figure 6.**
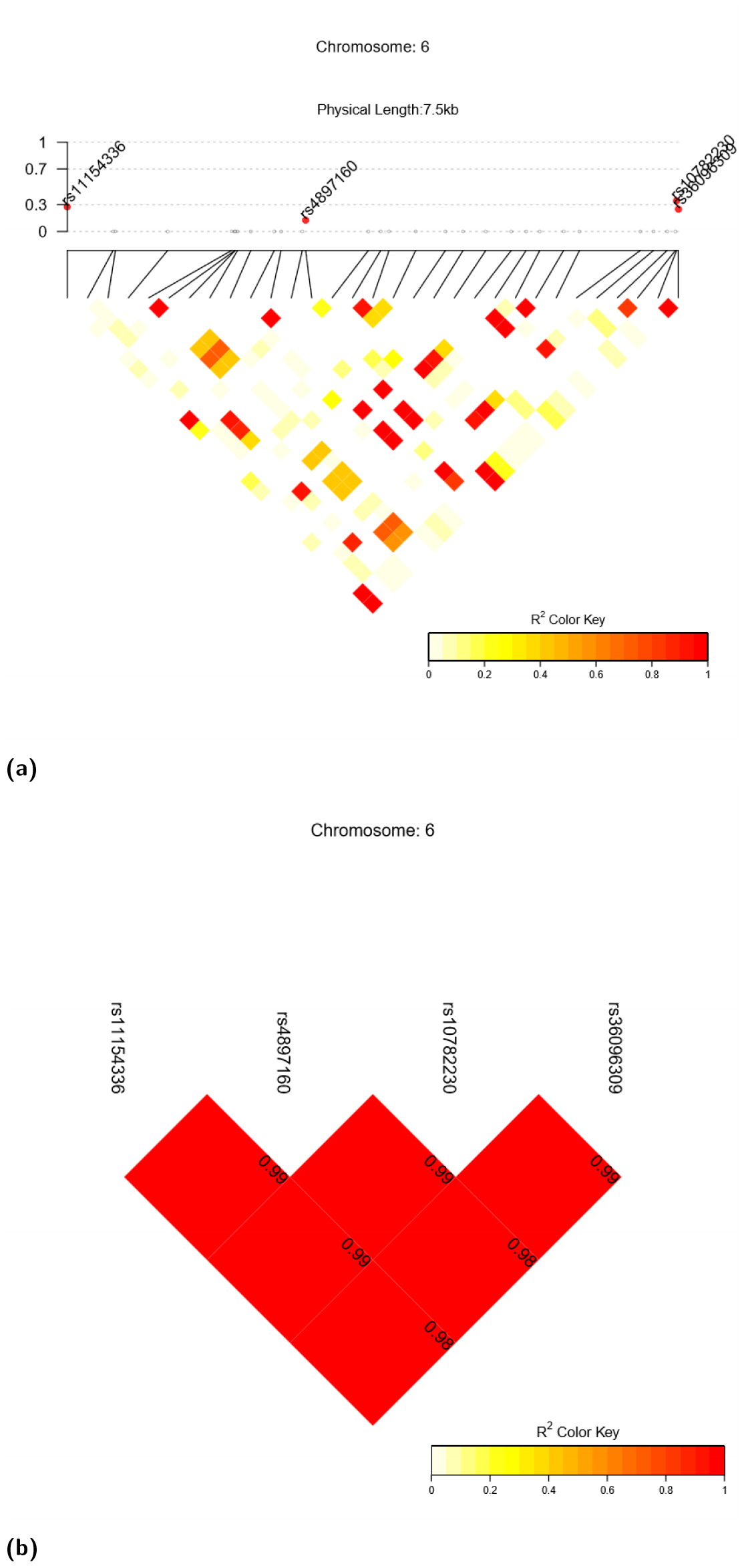
(a) SNP-specific Posterior Inclusion Probabilities (PIPs) for SNPs in a credible set identified using BHHT using a DS-BFDR threshold *α≤* 0.05. The triangle below the x-axis provides an LD heatmap among the SNPs. The solid red points indicate the SNPs included in the discovery sets identified through BHHT. (b) Correlation matrix between the SNPs included in the credible set discovered through BHHT.

## Discussion

In this study, we proposed a Bayesian hierarchical hypothesis testing (BHHT) procedure for identifying credible sets for problems involving highly collinear predictors. This setting is ubiquitous in modern GWAS studies where (owing to the continued decrease of sequencing costs) we are quickly progressing from using SNP arrays and imputed genotypes into whole-genome sequencing. Examples of this include the UK-Biobank (Sudlow *et al*. (2015) which has recently released whole-genome sequencing for nearly half a million individuals) and the All of Us program (All of Us Research Program (2019) which is on a path to releasing whole-genome sequencing data for hundreds of thousands of individuals from the US). The availability of whole-genome sequence genotypes will bring unprecedented opportunities for fine mapping of disease risk alleles because, at least in principle, fully sequenced genomes will enable the identification of causal variants without relying on LD between causal variants and those included in a genotyping array. However, whole-genome sequence-derived SNPs can be in extremely high LD, making fine mapping increasingly challenging. Analysis based on marginal association testing (e.g., single-SNP-phenotype association testing) will lead to many discoveries, but the vast majority of such discoveries will not be risk loci; instead, these will be SNPs in LD with risk variants.

To refine the results of marginal association tests, a common approach is to apply Bayesian variable selection methods to the SNP segments discovered using marginal association tests Benner *et al*. (2016). When this approach is used, risk loci are often identified using PIPs. However, as we show in this study, when LD is extremely high, the use of individual SNP PIPs can lead to a loss of power. This problem can be addressed using BHHT approach described in this study. BHHT can be used to identify individual SNPs with high PIP and credible sets, consisting of SNPs that are jointly confidently associated with the trait being studied. We have shown through simulations and real data analysis that these procedures can provide (1) high power with low FDR, (2) accurate error control, and (3) fine-mapping resolution. We further show that these approaches gain significant power over the tests for individual effects, which is the commonplace practice in GWAS.

In linear models, collinearity poses various challenges. Most popular variable selection procedures, e.g., lasso Tibshirani (1996), elastic net Zou and Hastie (2005), tend to produce a discovery set with many false discoveries when the features with non-zero effects are highly correlated with other features Su *et al*. (2017). Therefore, studies in this area have changed the objective from selecting individual predictors to credible set inference, which captures the uncertainty introduced by collinearity. In Bayesian variable selection, one approach (used in the SuSiE Wang *et al*. (2020) method) is to specify a prior distribution that leads to the identification of credible sets. Previous studies have shown that SuSiE outperforms the other procedures in the literature in terms of power, FDR, and size of credible sets (e.g., Wang *et al*. (2020), Sesia *et al*. (2020)). The problem of variable selection under collinearity has also been studied in the frequentist literature. Recent methods such as knockoffzoom Sesia *et al*. (2020) applied hierarchical testing to find the credible sets. Although our approach is similar there are important differences worth mentioning. First, the knockoffzoom procedure controls FDR at each resolution separately. Therefore, theoretically, it does not guarantee control over the discovery set FDR. In addition, knockoffzoom requires estimating the joint distribution of the inputs which, in many applications, can be challenging. This is not required in BHHT because in BHHT inferences are conditional on the observed data (including predictors). According to Sesia *et al*. (2020), knockoffzoom performs similar SuSiE (e.g., Figure 4 in Sesia *et al*. (2020)), the method we choose as a benchmark, in terms of power; however, SuSiE achieved higher resolution discoveries.

A potential advantage of the BHHT relative to SuSiE is that BHHT can be used with any variable selection prior. Therefore, as shown in our simulations, BHHT can outperform SuSiE if the assumptions made by the SuSiE prior do not hold (e.g. when there are many causal variants and extremely high LD). On the other hand, BHHT can sometimes produce very large credible sets–this is particularly a problem when one uses a very low FDR threshold (e.g., 0.01) which forces the method to include many SNPs in the set to meet such threshold. SuSiE does not have this limitation and hence in some cases, it can discover smaller credible sets. For this reason, in Fig. 3 when S=5, and r=0.90 we see that SuSiE shows a slight power gain over BHHT when the maximum credible set size was restricted to 5. However, when we allow larger credible sets in Fig. S3 the two methods exhibit similar power in the same simulation setting.

In our study, we focused on BHHT controlling the discovery-set BFDR (DS-BFDR). However, we also evaluated two more conservative FDR-control methods: (i) the node BFDR (i.e., controlling the BFDR for each credible set, as opposed to do it at the discovery-set level), and (ii) the subfamily-wise BFDR, a method proposed in a frequentist setting by Yekutieli (2008) and adapted by us to a Bayesian framework. Overall, the simulation and real-data analysis showed that all criteria performed similarly (see Fig. S4 and Fig. S6 in the Appendix). This may seem counter-intuitive because the node-FDR and the subfamily-wise BFDR are more conservative than the DS-BFDR. However, one reason for such a behavior is the distribution of the set-PIP values. We have observed that across clusters these probabilities exhibit a U-shaped distribution, with a high frequency of set-PIPs very close to zero for clusters not harboring causal variants and set-PIPs very close to one for those clusters harboring one or more causal variants. With such a distribution of the set-PIPs, BHHT has a very sharp transition from low to high FDR (regardless of the method used to control it); thus, the global and local control of the FDR leads to similar discovery sets.

There are many interesting avenues that can lead to further research in BHHT and credible set inference. Traditional hypothesis testing focuses on developing decision rules that maximize power controlling type-I error rate. However, in multi-resolution inferences one also needs to consider the mapping resolution achieved. That is, one needs to find a balance between power, type-I errors, and mapping resolution. In our study, we opted for comparing models by either limiting the credible set size (e.g., in simulation 1 where we followed Meinshausen (2008)) or by defining true and false discoveries based on mapping distance (e.g., in simulation 2 and in our real data analyses where we followed de los Campos *et al*. (2009)). However, a perhaps better approach would be to include the mapping resolution as one of the features of the decision making process. For instance, one could derive decision rules that minimize DS size subject to a BFDR control restrictions.

On the application side, the methods discussed in this study can be applicable to a wide range of fields beyond GWAS. An interesting application area is in the field of neuroscience, which focuses on identifying regions of the human brain related to neural disorders. Modern imaging techniques such as tractography Jeurissen *et al*. (2019) provide data on the individual neuronal tracts in the human brain, which has a highly complex correlation structure. The BHHT can be applied to these complex datasets to better understand the relationship between tracts and the disorder. In general, BHHT are suited to the area of image analysis. Since the images have structures where nearby pixels tend to be strongly correlated, one can apply the multi-resolution tests to identify a group of pixels exhibiting some properties of interest. Likewise, we believe there may be interesting applications of BHHT for analysis of high-dimensional phenotype/risk factors, where in this method could help identifying sets of correlated risk factors that are jointly associated with an outcome. Finally, in a GWAS setting, there are some potentially interesting extensions of the method proposed in this study for mapping sets of loci that are simultaneously associated with more than one trait or disease (i.e., for the study of pleiotropy).

In summary, the BHHT method proposed in this study can provide accurate and powerful inference in a wide range of scenarios. With the use of modern software and with access to computing clusters such as those available at many universities and research institutions, this method can be applied to ultra-high-dimensional datasets.

## Data availability

All data necessary for confirming the conclusions presented in the article are represented fully within the article. Genotype and phenotype data from the UK Biobank are available to all researchers upon application at http://www.ukbiobank.ac.uk/register-apply/. The scripts used to reproduce the results are available at https://github.com/AnirbanSamaddar/Bayes_HHT.

## Appendix

## Different Type-I errors

In the main paper, we described the discovery set BFDR (DS-BFDR) control. In this section, we describe two more conservative Type-I error rates in the hierarchical hypothesis testing framework and the algorithms to control them.

i. **Local (or node) BFDR (LFDR):** For each test, the local BFDR is the posterior probability of the null given the data: *LFDR*_Ω_ = 1 *−PIP*_Ω_ where Ω is the set of all coefficients under that node (see Eq. 5). Algorithm S1 describes the algorithm we use to traverse a tree, rejecting the null hypothesis provided that the *LFDR*_Ω_ at each node is smaller than a tolerable threshold (*α∈* [0, 1], e.g., *α* = 0.05). As the BFDR of a discovery set is the average of the LFDRs of its members, it follows that a decision rule that controls the LFDR for each of the elements of the DS yields DSs with a DS-BFDR *< α*. However, it is known that controlling the LFDRs of each element in the DS can be conservative.
ii. **Subfamily-wise BFDR:** A decision rule that controls the DS-BFDR does not bound the LFDR in each cluster. Hence, there can be cases where clusters with low inclusion probabilities are included in a DS not based on their merit but by benefiting from the fact that other clusters have very high inclusion probabilities. This behavior has been noted in multiple testing Siegmund *et al*. (2011) where many true rejections can inflate the denominator of the FDR so that it allows a small number of false discoveries. Therefore, following Yekutieli (2008), we consider a third procedure that controls the subfamily-wise FDR. This approach is more conservative than a method controlling the DS-BFDR and less conservative than those based on the node-wise BFDR in (i). Algorithm S4 describes an algorithm to control sub-familywise BFDR.

The results from BHHT discussed in the main paper were based on the DS-BFDR error control procedure. We also evaluated BHHT using the node (local) BFDR and the subfamilywise BFDR. Overall, there were no noticeable differences between the three error control methods (see Fig. S4 and Fig. S6 for results using the three methods for error control, for the same scenarios discussed in the simulation study section displayed in Fig. 3 and Fig. 5).

## Theoretical justification of FDR control

Consider the multiple regression problem in Eq. 1. We are interested in testing a set of *k* null hypotheses about the regression coefficients denoted by *H*_01_, …, *H*_0*K*_. These hypotheses can be simple for example *β*_0*j*_ : *β*_*j*_ = 0 vs *H*_1*j*_ : *β*_*j*_ ≠ 0, *∀j* = 1, …, *p* or can be composite for example *H*_0*j*_ : *β*_*j*_ = *β*_*j*_^*′*^ = 0 vs *H*_1*j*_ : at least one of *β*_*j*_ or 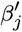 is not zero *∀j* = *j*^*′*^ = 1, …, *p*.

Let, *d*_*j*_ denote the decision rule for testing *H*_0*j*_ where *d*_*j*_ = 1 implies a rejection of *H*_0*j*_ and *d*_*j*_ = 0 implies otherwise. The FDR is defined by,

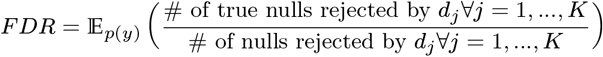

Note that the expectation is with respect to the marginal density p(y) which is not tractable. Therefore, following Genovese and Wasserman (2004), Müller *et al*. (2006), we use the Bayesian FDR or BFDR to estimate the FDR. BFDR is defined as,

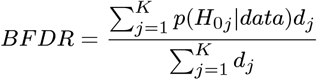

In the above definition, data represents the training data available that is *{y*_*i*_, *x*_*ij*_ : *i* = 1, …, *n*; *j* = 1, …, *p}*. By the below lemma, we show that the BFDR is an unbiased estimate of the FDR (over conceptual repeated sampling from *p*(*y*)). Therefore, by controlling the BFDR we control the FDR at a desired level. The proof is a straightforward application of the law of the iterated expectation.

**Lemma S1**. *Under the above set up, FDR* = E_*p*(*y*)_(*BFDR*)

*Proof*. From the definition,

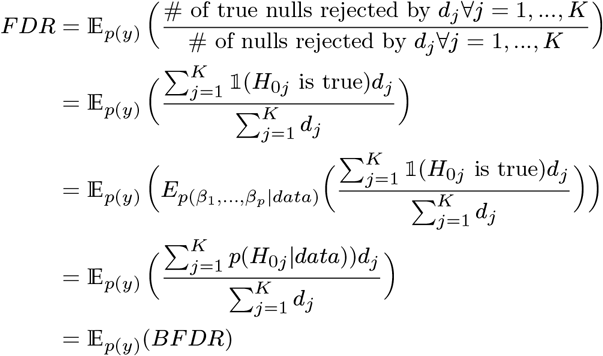

The third equality is due to the law of iterated expectation. The fourth equality is valid since *d*_*j*_ *∀j* = 1, …, *K* are functions of the training data and therefore we can apply the conditional expectation only to the indicators 𝟙_(*H*_0*j*_ is true) *∀ j* = 1, …, *K* in the numerator. Hence the proof.

## Set up for the algorithms

Let, m hierarchical hypotheses in a tree be denoted by (*H*_01_, *H*_11_), …, (*H*_0*m*_, *H*_1*m*_), where (*H*_0*j*_, *H*_1*j*_) denotes the null (i.e., none of the predictors in the cluster has an effect) and the alternative (at least one of the predictors belonging to the node has an effect) in the j-th cluster. Following Eq. 5, the posterior samples can be used to estimate the set-PIPs at each cluster denoted by, *v*_*j*_ = *p*(*H*_1*j*_ *data*) *∀j* = 1, …, *m*.

## Node BFDR control

We describe the node BFDR control algorithm –

### Algorithm S1

Node BFDR control

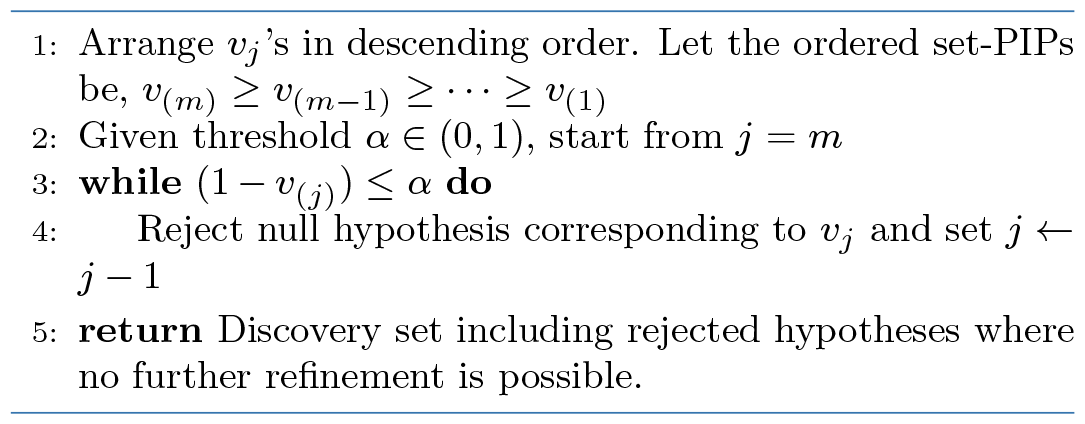

## DS-BFDR control

For the DS-BFDR control, we control the DS-BFDR for the discovery set selected by Algorithm S1 for decreasing thresholds given by the ordered set-PIPs *v*_(*m*)_, *v*_(*m−*1)_, …, *v*_(1)_. The algorithm is as follows –

### Algorithm S2

DS-BFDR control

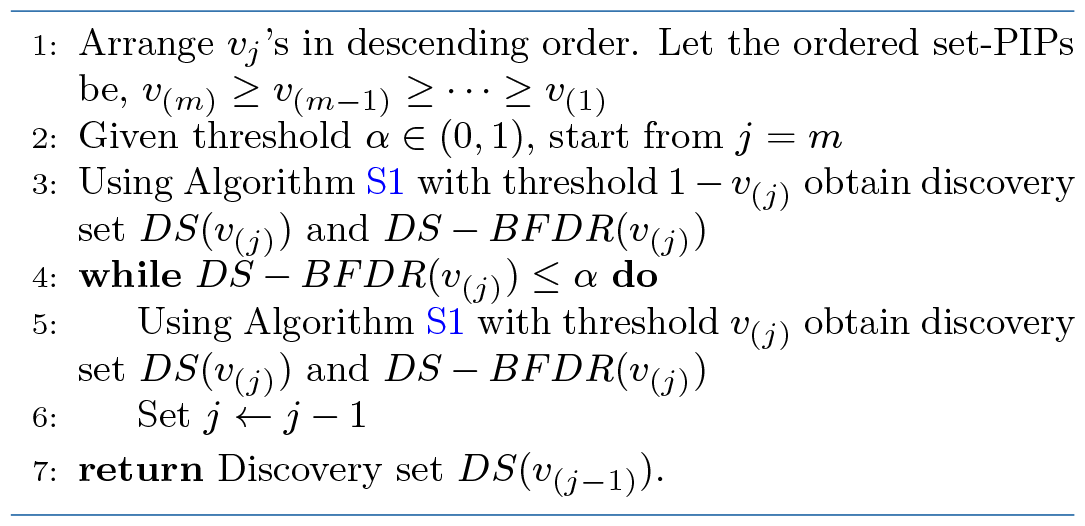

**Marginal hypothesis testing:** Consider the multiple regression problem in Eq. 1. We are interested in testing a set of p marginal null hypotheses denoted by *H*_01_, …, *H*_0*p*_ where *H*_0*j*_ : *β*_*j*_ = 0 vs *H*_1*j*_ : *β*_*j*_ ≠ 0, *∀j* = 1, …, *p*. Given the PIPs denoted by *v*_*j*_ *j* = 1, …, *p* and a threshold *α* ∈ (0, 1), we outline the below steps for marginal hypothesis testing with BFDR controlled at a desired level –

### Algorithm S3

Marginal hypothesis testing

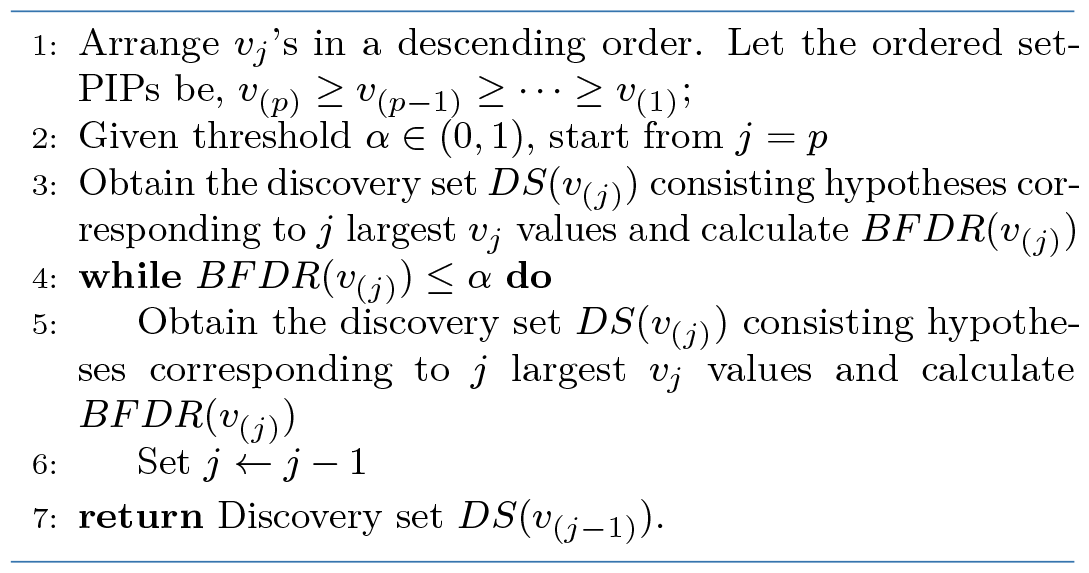

## Subfamily-wise FDR control

In a hierarchy, a sub-family is defined by the clusters that share the same parent. For example, in Fig. S1, the parent node hypothesis is *H*_0_ : *β*_1_ = *β*_2_ = *β*_3_ = *β*_4_ = 0 Vs *H*_0_: at least one coeff. ≠ 0 which is marked by solid red squares. The subfamily of the parent node is marked by solid blue rectangles. The subfamily-wise FDR is defined as the FDR in the discovery set within a subfamily. Therefore, to control the subfamily-wise FDR we use Algorithm S3 for each subfamily starting from the top node. The algorithm is described below –

### Algorithm S4

Subfamily-wise BFDR control

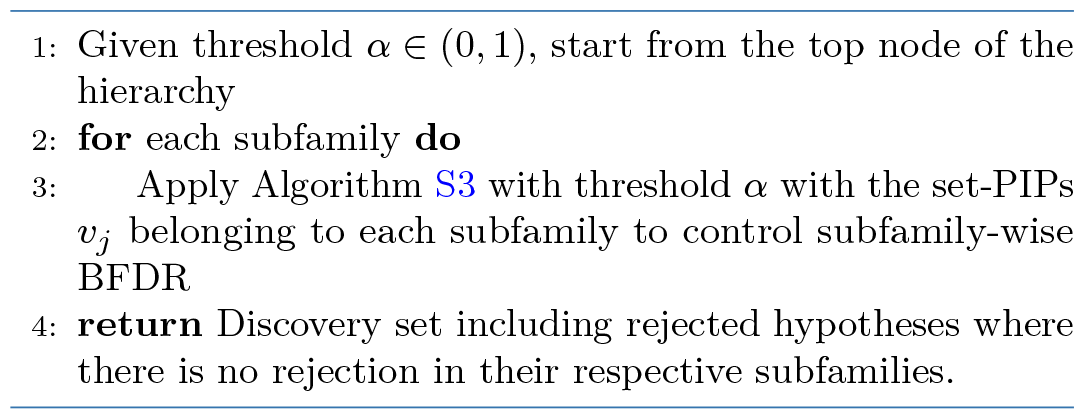

**Figure S1.**
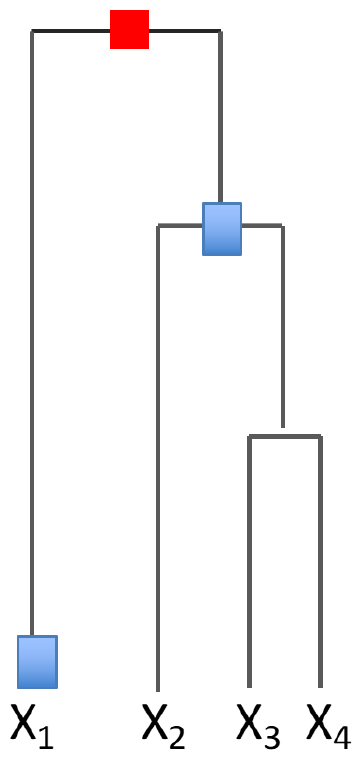
Demonstrating subfamily

## Supplementary Data

### Generation of the simulation study covariates

In this section, we provide additional details regarding the generation of the covariates *{x*_*ij*_ : *i* = 1, …, *n*; *j* = 1, …, *p}* for the simulation study. As mentioned in the Simulation study section, we have generated covariates where *Cor*(*x*_*i*(*j*)_, *x*_*i*(*j*+*k*)_) = *ρ*^*k*^ decays with distance *k*. Here, *i* is an index for the subject, *j* is an index for the feature in the sequence, and *k* is the lag-between predictors. To generate the columns of the covariate matrix in this way we followed the below two steps,

1. Generate the first column of the covariate matrix with the n elements from the i.i.d. *Binomial*(2, *a*). We set *a* = 0.2 in all simulations.
2. Generate the j-th column by randomly permuting a random fraction (***f***) of elements of (j-1)-th column. The random fraction ***f ∼ beta*(*a, b*)**.

By changing the values of (*a, b*), we simulate different degrees of correlation between adjacent columns. For example, setting (*a, b*) = (2, 3) implies correlation *∼*0.45 and (*a, b*) = (0.5, 3) implies correlation *∼*0.9. Note that, as the distance between two columns grows, the correlation decreases as more elements change positions randomly due to permutation.

### Model hyper-parameters

In this section, we provide details of the hyperparameters used in the simulation study and the real data application. In this study, we used two Bayesian models to fit the data, which are independent spike-and-slab regression and SuSiE. We have used the respective R-packages BGLR Pérez and de los Campos (2014) and susieR Wang *et al*. (2020) to fit the models. Below are the models and key hyperparameter values used.

**Table S1.**
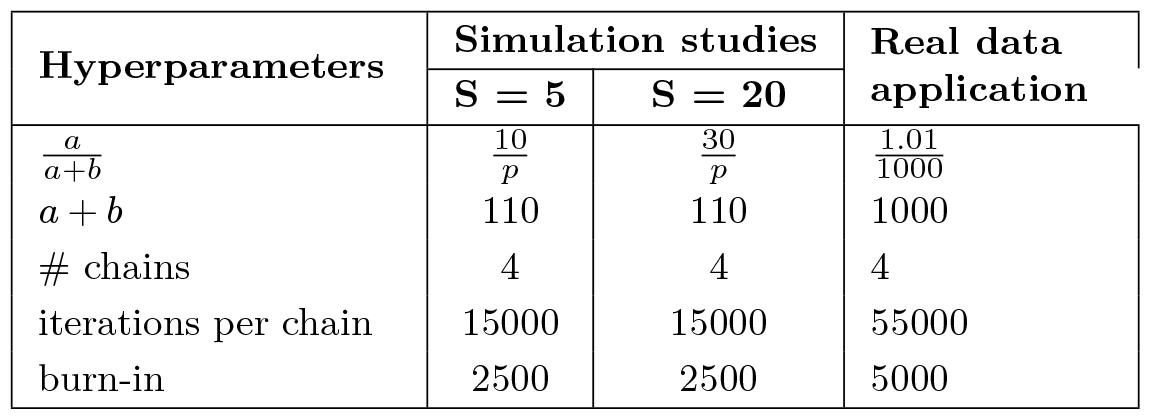
Table showing hyperparameter values for simulation study and real data application for the spike-and-slab model.

### Independent spike-and-slab model

The Bayesian spike- and-slab model (aka BayesC) puts a point mass and a gaussian slab on the regression coefficients. The model hierarchy is as follows,

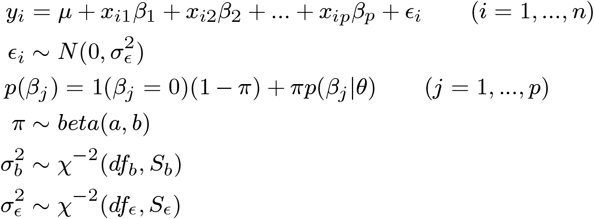

Here, *χ*^*−*2^ denotes the scaled-inverse chi-square distribution. Also, since we used the Gibbs sampler to draw samples from the posterior, a few other important hyperparameters are the number of chains, number of iterations, and number of burn-in steps. We set the hyperparameters according to Table S1. Note that the only hyperparameter changed between the two simulation settings, *S* = 5 and *S* = 10, is the prior probability of inclusion *a/*(*a* + *b*). In our simulations, if we hold this to be fixed at 10*/p* for *S* = 20 the power of the BHHT method drops marginally. However, the key reason to change this hyperparameter with *S* is to be consistent with SuSiE (described below). For the other parameters, we set them to the default values of the software implementation.

### The sum of Single Effects model (SuSiE)

The SuSiE model assumes that out of the p coefficients L of them are non-zero. The model hierarchy (as given in Wang *et al*. (2020)) is as follows,

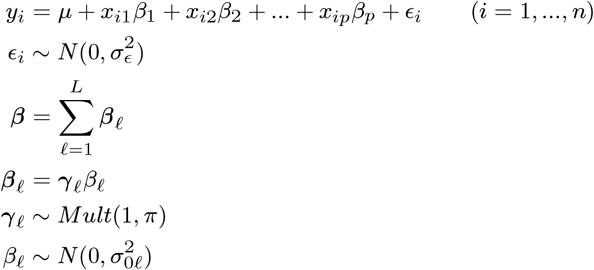

In the above set of equations, ***β*** represents the *p*×1 dimensional vector of regression coefficients. The hyperparameter *L* here has a similar interpretation as the prior probability of inclusion in the spike-and-slab model. However, the misspecification of *L* has a significant impact on the power of SuSiE.

Especially when *L << S*, the SuSiE method tends to lose significant power. The general advice in Wang *et al*. (2020) is to choose *L* to be higher than *S*. Therefore, for the simulation settings where *S* = 5 and *S* = 10 we fix *L* = 10 and *L* = 30, respectively, to be consistent with the spike-and-slab prior. The other hyperparameters in the model are set to the default values of the software implementation.

### Real data modeling strategy

In this section, we discuss additional details used to fit a Bayesian linear regression model using the calls (*∼*785,000 SNPs) and imputed (*∼*15 million SNPs) datasets. Previously we discussed the modelling strategy on the imputed dataset. For calls, following Funkhouser *et al*. (2020) we created windows of 2900 SNPs by taking disjoint cores of 1500 SNPs and adding overlapping flanking regions of 700 SNPs on both sides. We fit the Bayesian model on the entire window (2900 SNPs) however only retrieve the samples from the core (1500 SNPs).

We collected samples from the posterior distribution by using the BLRXy function of the BGLR R-package Pérez and de los Campos (2014). This function implements the same set of models implemented in BGLR but performs the computation using sufficient statistics (*X*^*′*^*X, X*^*′*^*y, y*^*′*^*y*); where *X* is the design matrix and y is the response vector. Following this strategy offers great computational advantages when *n >> p*.

## Supplementary figures

Below are the supplementary figures that are referred to in the main paper.

**Figure S2.**
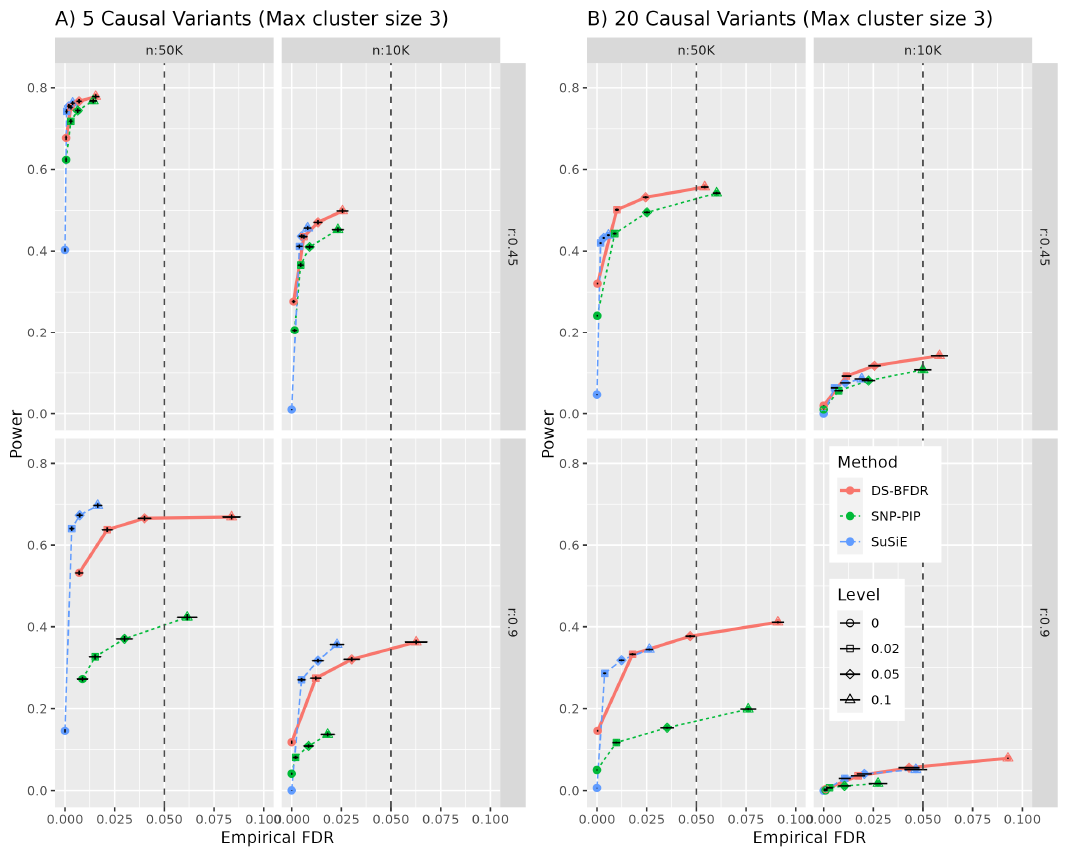
Power vs FDR curves for <u>simulation one</u>, by restricting maximum cluster size to 3. Here, *α, n*, and *r* indicate the BFDR threshold used, sample size, and the lag-1 correlation of the simulated predictors, respectively. The Left and right panel represents scenarios where there are 5 and 20 causal variants respectively. We hold *h*^2^ fixed at 1.25% for all settings. Putting more restriction on the cluster size criteria have reduced the power advantage of CS approaches relative to marginal SNP-PIP inference.

**Figure S3.**
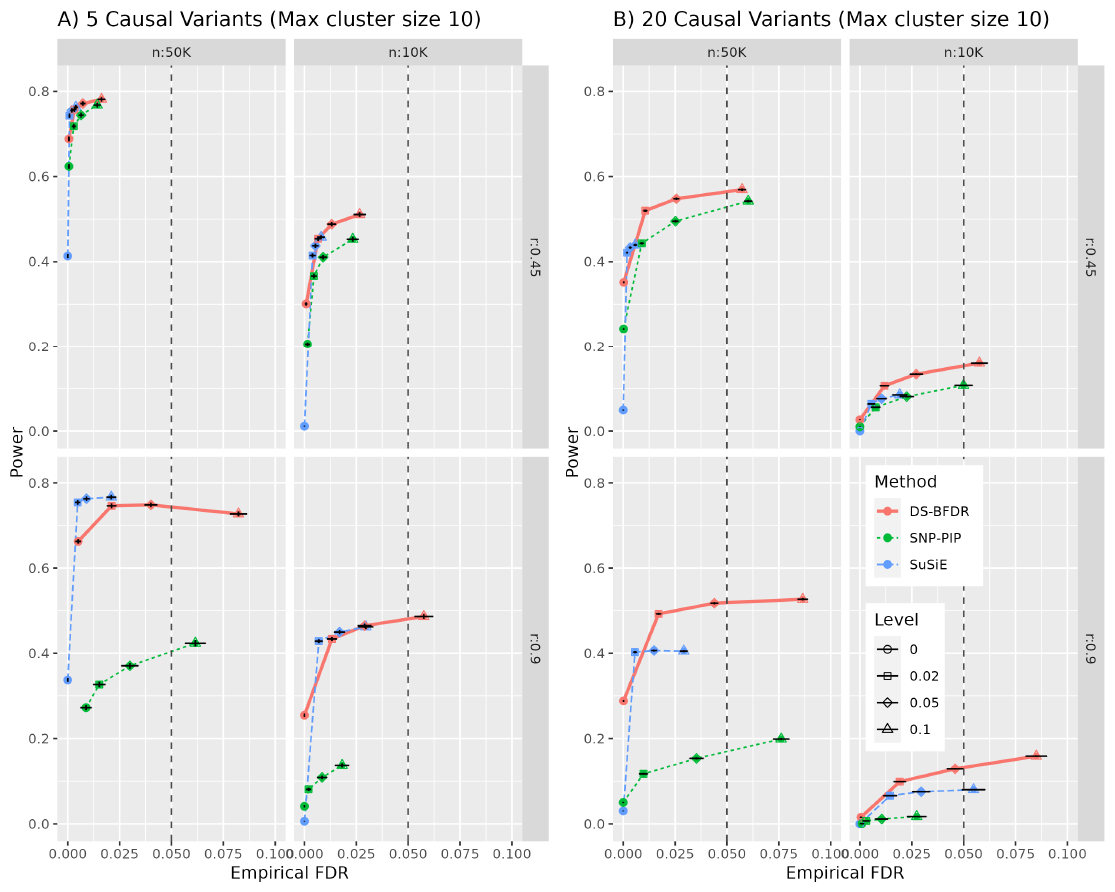
Power vs FDR curves for <u>simulation one</u>, by restricting maximum cluster size to 10. Here, *α, n*, and *r* indicate the BFDR threshold used, sample size, and the lag-1 correlation of the simulated predictors, respectively. The Left and right panel represents scenarios where there are 5 and 20 causal variants respectively. We hold *h*^2^ fixed at 1.25% for all settings. Putting less restriction on the cluster size criteria has increased the power advantage of CS approaches relative to marginal SNP-PIP inference. Furthermore, the DS-BFDR approach gains more power due to lower-resolution discoveries relative to SuSiE.

**Figure S4.**
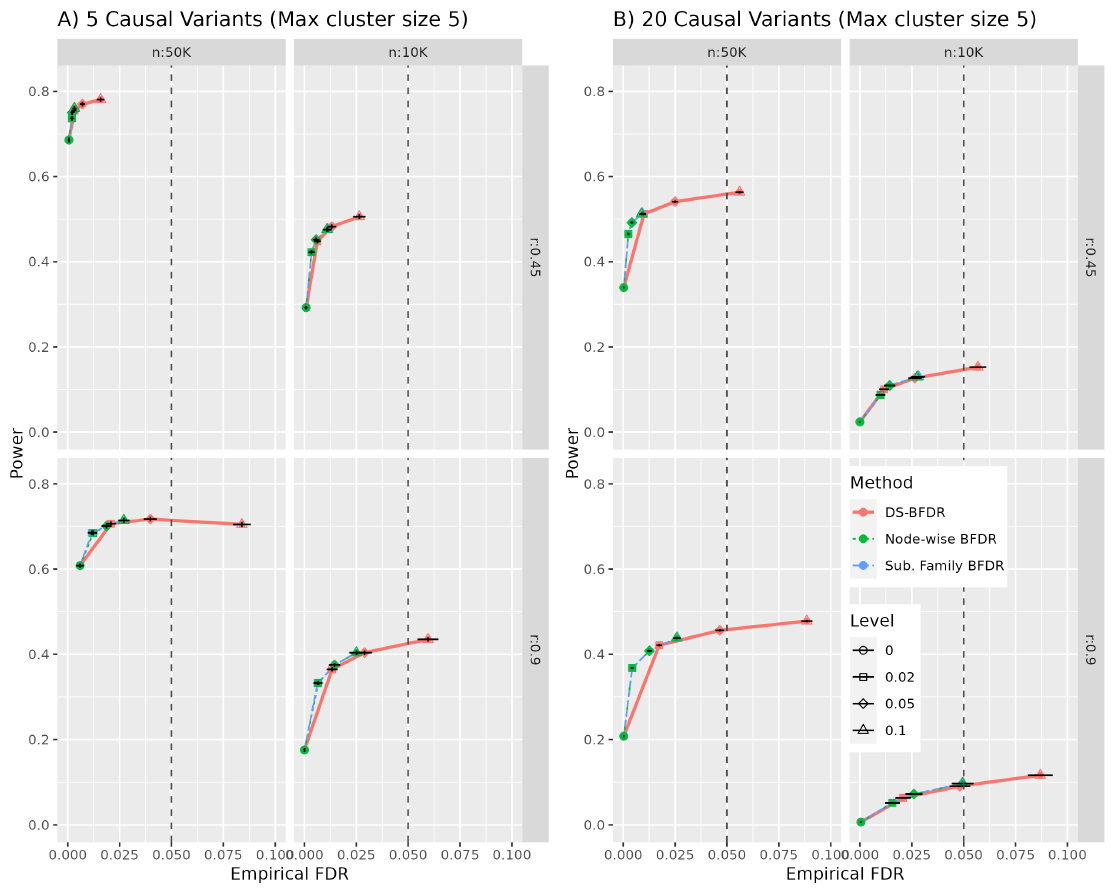
Power vs FDR curves of the three error controlling approaches for <u>simulation one</u>. Here, *α, n*, and *r* indicate the BFDR threshold used, sample size, and the lag-1 correlation of the simulated predictors, respectively. The Left and right panel represents scenarios where there are 5 and 20 causal variants respectively. We hold *h*^2^ fixed at 1.25% for all settings. All the error-controlling approaches show similar performance.

**Figure S5.**
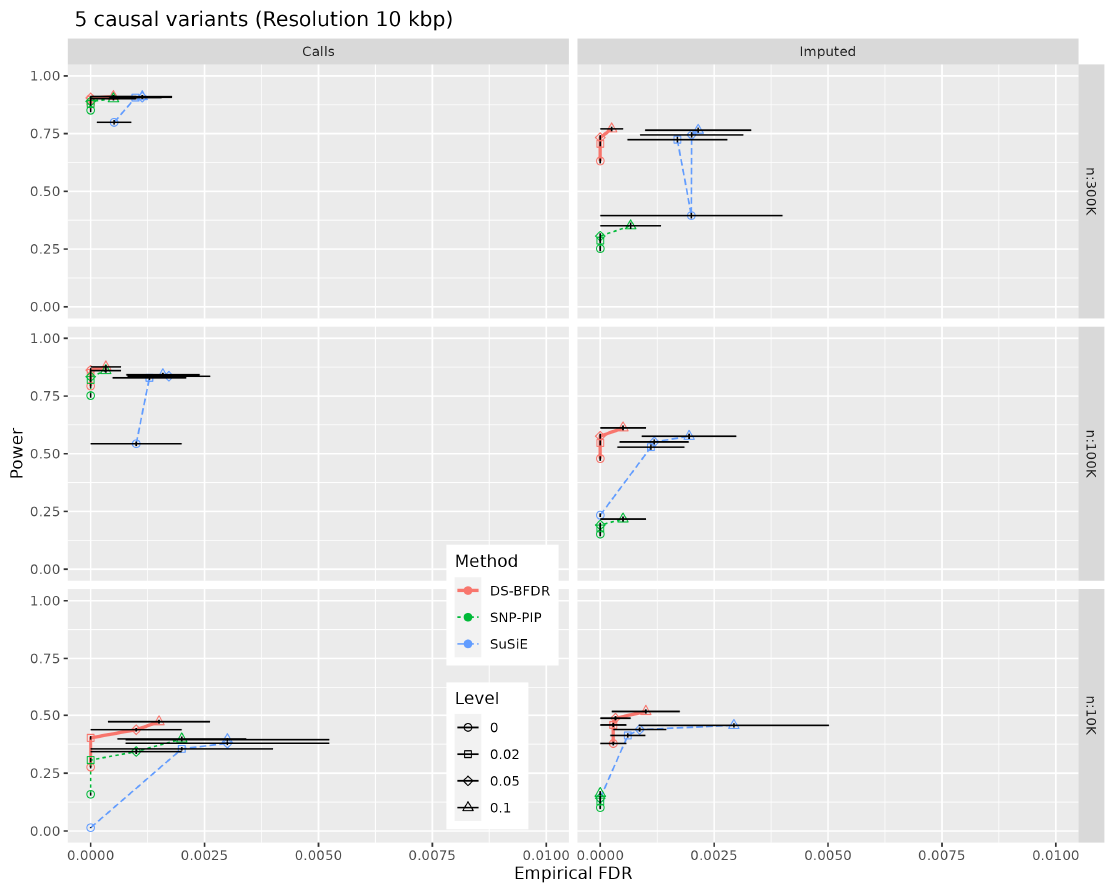
Power vs FDR curves for simulation two by sample size (rows) and the SNP set used (columns) at a low resolution of 10 kbp. Here, *α* and *n* indicate the BFDR threshold used and sample size respectively. In all scenarios, SNPs explained 1.25% of the variance of the outcome. As we lower the resolution from 0 in Fig. 5 to 10 kbp all methods have significantly less FDR while maintaining power at similar levels.

**Figure S6.**
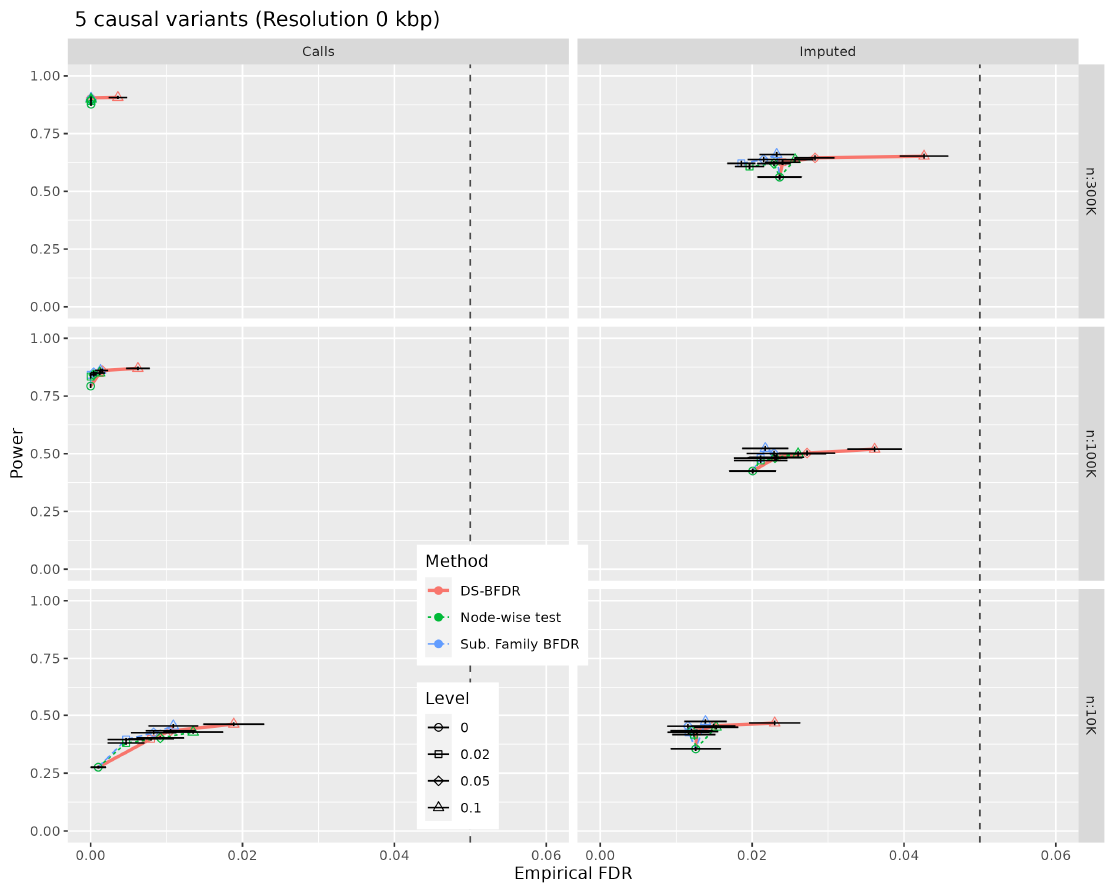
Power vs FDR curves for simulation two by sample size (rows) and the SNP set used (columns) for different error control strategies. Here, *α* and *n* indicate the BFDR threshold used and sample size respectively. In all scenarios, SNPs explained 1.25% of the variance of the outcome. There are no noticeable differences between the three strategies in terms of power however DS-BFDR error control provides a tighter control over FDR to the desired level.

**Figure S7.**
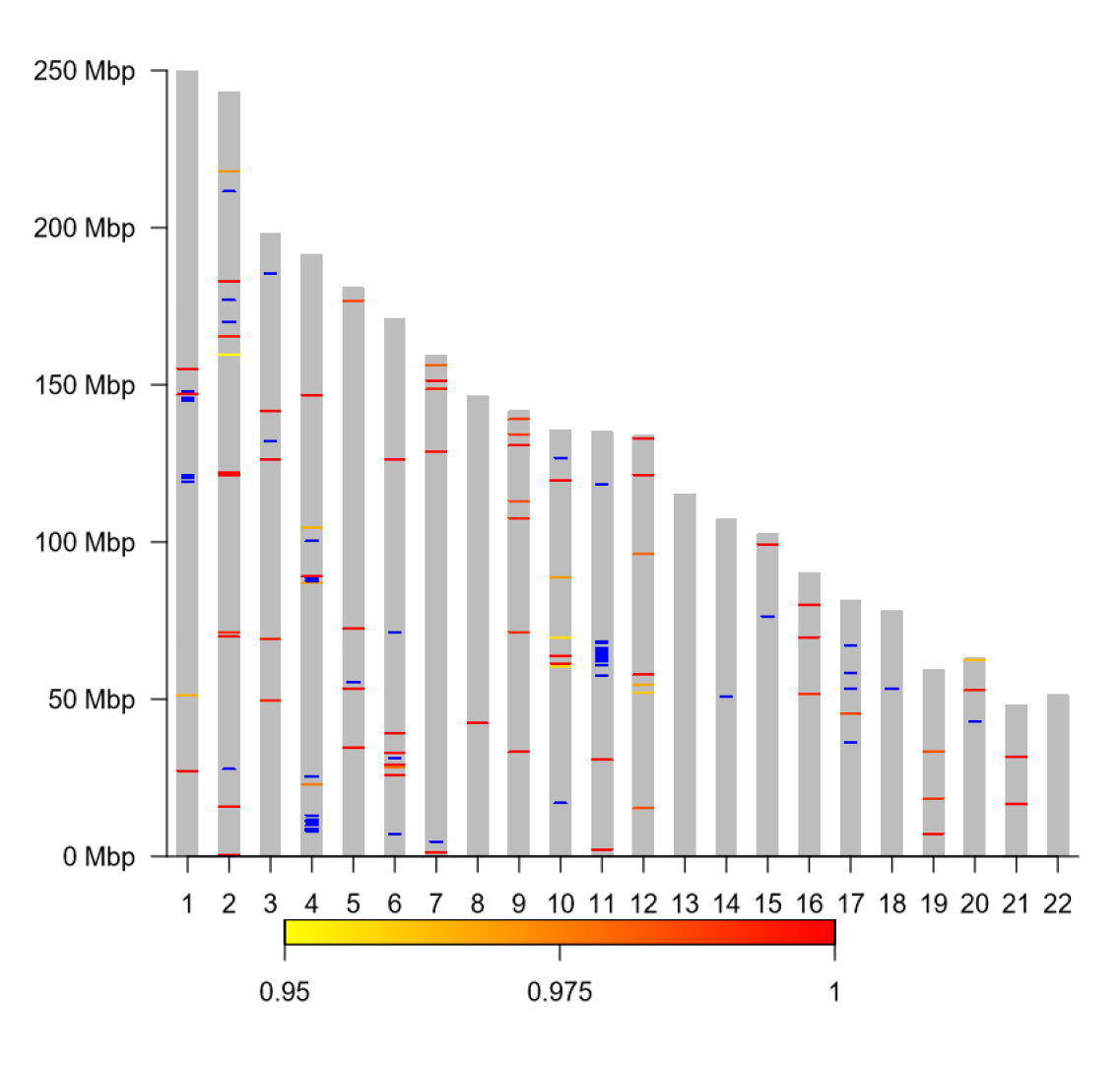
Ideogram displaying each of the 22 autosomal chromosomes for the imputed dataset and singleton and segment discovered in the GWAS of the trait serum urate. The individual SNPs that cleared the 0.05 BFDR threshold are represented using blue lines along with the segments identified which are represented by bars colored as per their joint probability of inclusion in a yellow-to-red scale.

